# Decreased Damage for proton FLASH vs Conventional Dose Rates in Mouse Jejunum Shown by Quantitative Assessment of γ-H2AX

**DOI:** 10.64898/2026.09.11.750404

**Authors:** Najah Curtis, Michele M. Kim, Ioannis Verginadis, Wei Zou, Eric S. Diffenderfer, Constantinos Koumenis, Cameron J. Koch, Rodney D. Wiersma

**Affiliations:** University of Pennsylvania; University of California, Los Angeles

## Abstract

**Purpose:** FLASH radiation with ultra-high dose rate delivery is less damaging to normal tissue than conventional radiation ( <1 Gy/s). Since radiation depletes oxygen (ROD), this damage reduction might occur via the oxygen effect. ROD experiments have shown an oxygen-independent reduction in dose effectiveness at FLASH dose rates. However, prior in vivo ROD measurements relied on extracellular oxygen probes that could not penetrate cell membranes, leaving intracellular effects unresolved. To investigate the ROD hypothesis more directly, we developed a novel three-component immunohistochemical assay with algorithmic image processing to quantitatively compare DNA damage following FLASH and conventional irradiation in mouse jejunum.

**Methods:** Mice received intravenous EF5 2 hours before proton irradiation at FLASH (103.63 +/- 17.2 Gy/s) or conventional (0.73 +/- 0.1 Gy/s) dose rates of 2.5 Gy or 5 Gy, with unirradiated controls. Mice were euthanized 30 minutes post-irradiation, and 10 cm of jejunum was frozen as a ‘Swiss Roll’, sectioned, stained, and imaged. Tissue sections were stained for γ-H2AX, DRAQ5, and EF5 to assess DNA double-strand breaks, total DNA content, and hypoxia, respectively. An in-house algorithm identified individual cell nuclei and registered each nucleus with its corresponding γ-H2AX and EF5 signals, enabling quantitative measurement of DNA damage as a function of local tissue hypoxia.

**Results:** Hypoxia was greatest in the villi and, to a lesser extent, the outer jejunal musculature, with substantial inter-animal variation. DNA damage decreased in hypoxic regions. FLASH enhanced the hypoxia-associated reduction in DNA damage compared with conventional dose rate and, separately, revealed an oxygen-independent reduction in DNA damage, suggesting an additional FLASH sparing mechanism.

**Conclusion:** Current results suggest that FLASH compared to conventional dose rate radiation caused less DNA damage with increasing effect at low oxygen levels, a result consistent with ROD as a mechanism. Pronounced tissue heterogeneity in murine jejunum requires further studies to segment the effect for each tissue type.

## Introduction

FLASH radiotherapy delivered with ultra-high dose rate is currently being investigated for its potential for decreased normal tissue damage compared to conventional radiation (<1 Gy/s). Since FLASH spares normal tissue without reducing tumor control, it increases the therapeutic ratio^1–6^.

An initial hypothesis for the mechanism of the FLASH radioprotective effect was radiation-chemical oxygen depletion (ROD). Hypoxic cells are ∼3-fold more resistant than aerobic cells, with half maximal resistance *in vitro* at ∼5 mm Hg ^7^. Thus, if cells critical for tissue survival exist near this half-maximal value, depletion of oxygen via ROD during the FLASH dose would decrease the local tissue pO_2_, and cause additional radiation resistance leading to protection^8,9^. At conventional dose rates, a similar amount of oxygen would be consumed for the same total dose but the local tissue pO_2_ would be maintained due to rapid resupply by the tissue vasculature^10^. Thus, the ROD hypothesis for the FLASH mechanism requires a sufficient ROD at the doses used, and also a pre-irradiation level of oxygenation in the appropriate range. Furthermore, since tumors are already hypoxic, they may not experience additional protection from FLASH.

A second possible mechanism for FLASH radioprotection, which we refer to here as the normal tissue protection factor (FLASH-ntPF), has emerged from ROD experiments. Using albumin solutions, the *‘G’* value (µM/Gy) for ROD was lower at FLASH than conventional dose rates, indicating a direct radiation-chemical impact. This mechanism may be related to free radical lifetimes, an enhanced role of cellular antioxidants, or possibly the radiation chemical depletion/production of other species at FLASH dose rates^11–15^.

To date, there is no research demonstrating a relationship between ROD measurements in protein solutions and DNA damage in normal tissue, even though differences in DNA damage would be the most likely cause of changes in radioresistance. Therefore, a more informative mechanistic endpoint for evaluating the FLASH mechanism *in vivo* is the direct, quantitative measurement of DNA damage in intact tissue combined with an appropriate measure of tissue pO_2_.

To allow such measurements, we have developed a novel three-component immunohistochemical (IHC) assay that uses pixel-by-pixel image processing to quantify tissue hypoxia, DNA content, and DNA damage at subcellular resolution in response to FLASH and conventional radiation. This approach allows the biological consequence most directly relevant to cellular toxicity — radiation-induced DNA damage — to be measured in intact tissue post-irradiation. This work thus presents the first study directly investigating DNA damage using mouse jejunum, a tissue known to exhibit the FLASH effect^2,4^, and to assess if any observed reduction in DNA damage in FLASH irradiated tissues can be attributed to either of the two mechanisms described above.

## Methods and Materials

### Experimental approach

The three-component assay was developed to quantitatively relate radiation-induced DNA damage to local tissue hypoxia within intact jejunum. The overall experimental and image-analysis workflow is summarized in **Fig. 1**. EF5, a hypoxia-sensitive 2-nitroimidazole, was administered 2.5 hours before irradiation to provide a spatial measure of pre-treatment tissue hypoxia. Mice were subsequently irradiated at FLASH or conventional dose rates and euthanized 30 minutes later to allow development of the γ-H2AX DNA damage signal (**Fig. 1a**). Jejunum was collected as a Swiss roll, sectioned, and stained for three complementary markers: γ-H2AX (GH) for radiation-induced DNA double-strand breaks, EF5 for tissue hypoxia, and DRAQ5 (DQ) for total DNA content (**Fig. 1b**).

**Figure 1.**
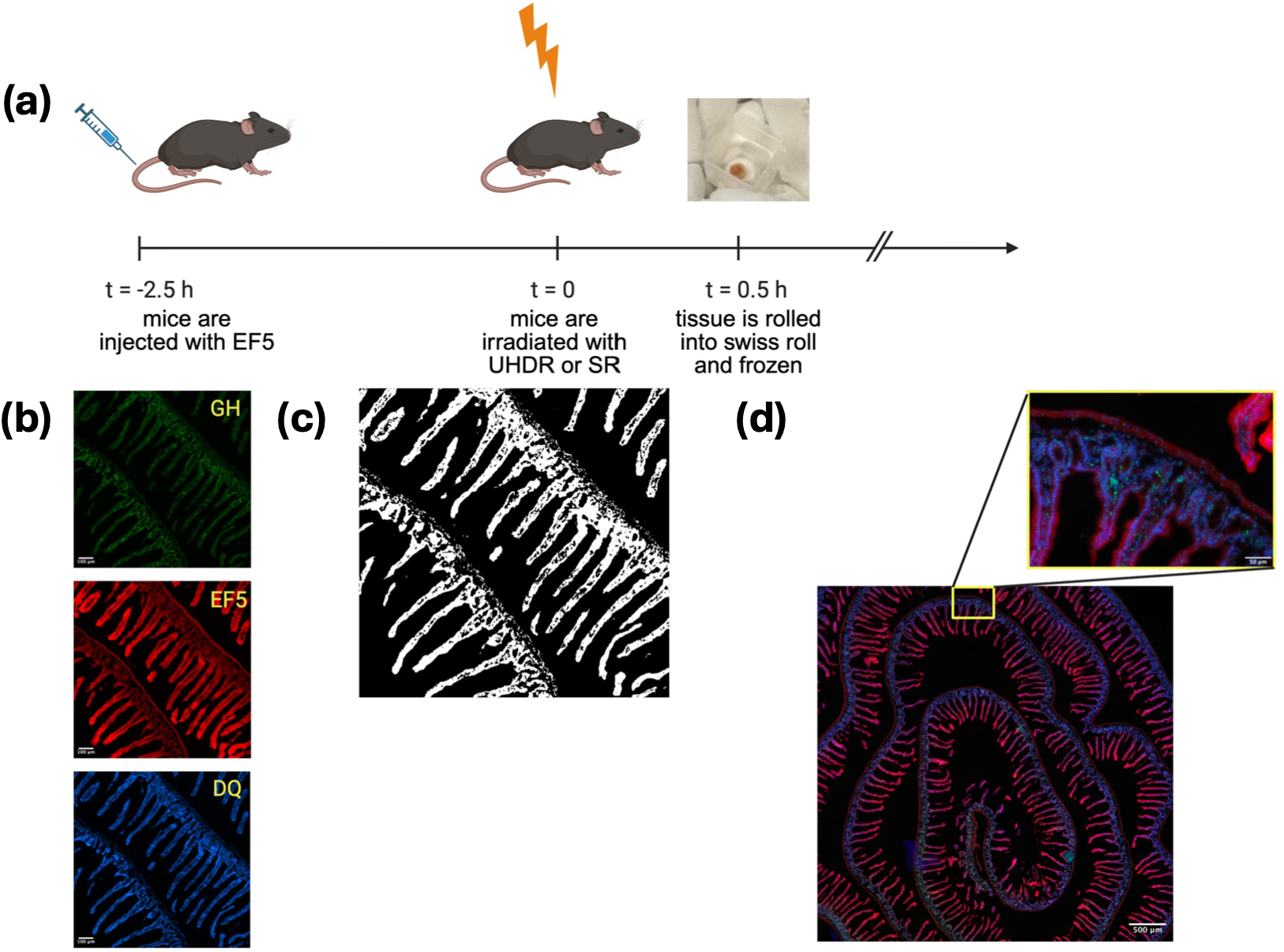
Overview of the quantitative three-component immunohistochemical method. (a) Shows the timeline of injection, irradiation, and tissue collection. Mice were injected with EF5 2.5 hours prior to irradiation, then tissue was collected 30 minutes post-irradiation. (b) Shows representative images for the γ-H2AX (GH), EF5, and DRAQ5 (DQ) stains. (c) shows the binary threshold mask created by the computational program that separates the tissue from the background. (d) Shows the mosaic composite of all three stains and the heterogeneity of EF5 throughout the jejunum. Image was created with BioRender.com.

For each tissue section, separate γ-H2AX, EF5, and DRAQ5 fluorescence mosaics were acquired sequentially using a precision automated microscope stage. An in-house algorithm then combined the DRAQ5 and γ-H2AX images to generate a binary analysis mask that segmented tissue-containing pixels from background (**Fig. 1c**). For every pixel included in the mask, the algorithm used its common X–Y position to extract the corresponding DRAQ5, γ-H2AX, and EF5 fluorescence intensities. γ-H2AX intensity was normalized to DRAQ5 intensity to account for differences in DNA content, producing a normalized DNA-damage measurement (GH_n_) for each analyzed pixel. Each GH_n_ value could therefore be directly associated with the EF5-derived hypoxia measurement at the corresponding tissue location.

Sequential microscope fields were assembled into large three-channel mosaics to sample the substantial spatial heterogeneity of the jejunum (**Fig. 1d**). The resulting pixel-level dataset allowed DNA damage to be quantitatively evaluated as a function of local tissue hypoxia, radiation dose, and dose rate. In particular, the relationship between normalized γ-H2AX and EF5 was used to determine whether increasing pre-treatment hypoxia was associated with reduced DNA damage and whether this relationship was enhanced following FLASH irradiation, as predicted by the ROD hypothesis. Detailed procedures for image acquisition, mosaic construction, fluorescence correction, masking, and quantitative analysis are described in the Supplement.

### Proton Irradiation

This study used the proton beam setup as described by Diffenderfer *et al*.^2^. IBA Proteus Plus (Louvain-La-Neuve, Belgium) delivered protons at 230 MeV (range ∼32.0 g/cm2) in the dedicated research room^16^. The protons were delivered in 2 ns pulses at 106 MHz, a temporal structure that is considered to be a continuous source. Alignment was performed using external horizontal and vertical lasers. Dose and dose rate were measured at the isocenter prior to each experiment using a NIST-traceable parallel plate Advanced Markus Chamber (PTW, Freiburg, Germany). Dose monitoring was done for each mouse with an online secondary parallel plate ionization chamber that was cross calibrated with the Markus Chamber at the treatment isocenter^17^. A 2-cm square collimator allowed whole abdomen irradiation.

### *In vivo* murine experiments

Nine-week old female C57BL6 mice (The Jackson Laboratory) were housed in AALAC-accredited facilities. All procedures were approved by IACUC. In order to determine the level of hypoxia in the tissue, EF5 was administered 2 hours prior to radiation (0.03 mg/g). EF5 is largely metabolized and/or excreted during this time, so the EF5 signal will detect the average level of tissue hypoxia before radiation^18^. Mice were irradiated with 5 Gy (n=3) or 2.5 Gy (n=3) at FLASH (103.63 +/- 17.2 Gy/s) or conventional (0.73 +/- 0.1 Gy/s) dose rates. For each experiment, each mouse received FLASH, conventional, or no (control) irradiation.

Mice were anesthetized during all irradiations using 1.5%-2% isoflurane in air supplied through a nose cone, then immediately allowed to recover. The mice were positioned such that the beam isocenter was 2cm superior from the hip bone, and 0.8cm above the mouse stage.

### Tissue Collection

At 30’ post-irradiation, each mouse was euthanized for tissue collection of its jejunum. The jejunum was extracted and rolled into a Swiss roll as introduced by Moolenbeek & Ruitenberg^19^. Approximately 10 centimeters of the jejunum was rinsed with PBS. The inside of the intestine was flushed with PBS to remove feces without damaging the intestine, then flushed with diluted OCT (80% OCT, 20% distilled water) and carefully cut longitudinally. The jejunum was rolled from inferior to superior using forceps. The Swiss roll was prevented from unraveling using 3M VetBond Tissue Adhesive on the last half centimeter of tissue to be rolled. It was immediately placed in a polyethylene mold filled with Tissue-Tek OCT, then frozen using dry-ice surrounding the molds.

### Fixation

Tissue sections were fixed, blocked, stained and rinsed as described by Koch^20^. 10-μm-thick sections (Leica Cryostat, Nussloch, Germany) adhered to Newcomer Supply Poly-L-Lysine Adhesive Microscope Slides (5010, 5013) were immediately fixed in graded ethanols (70% for 30’, 85% for 10’, 100% for 5’, 70% for 5’) then immediately placed in phosphate buffered saline with 0.3% Tween^®^ 20 (PBStw20), all at 4°C. Slides were placed in PBStw20 for hydration for about 1 minute before the area surrounding tissue was dried in order to ring the tissue using a Liquid Blocker Super PAP Pen as close to the tissue as possible while still leaving a slight amount of room for wicking off the block and stain.

### Immunofluorescence Staining

Two solutions were used in addition to PBS and PBStw20: Ab Carrier (PBStw20 + 15 mg/ml albumin, and Block (Ab Carrier with 2.5% (g/ml) fat-free milk powder & 5% (ml/ml mouse serum). Block solution was added to each slide for 5 hrs at 4°C.

Alexa Fluor^®^ 488-conjugated antibody (clone JBW301; MilliporeSigma, Burlington, MA) and Cy3-conjugated ELK-51 (made by Koch) were mixed with 2/3 antibody carrier and 1/3 block, resulting in a final antibody concentration of 50 μg/ml for anti-EF5 and 7 μg/ml for anti-γ-H2AX. The tissues were stained overnight at 4°C.

After staining, the sections were rinsed at 4°C: 2×30’ in PBStw20 and 1×30’ in PBS, then stored in PBS with 2% paraformaldehyde (pF) overnight at 4°. This fixes all the antibodies in place without affecting the fluorophores^20^.

Just prior to imaging, each slide was rinsed in PBS (RT), and DNA was stained using 10µM DRAQ5 (DQ) for 20’ at RT – see Supplement Methods. This was removed, slide rinsed, and a hemocytometer coverslip with 100 µm thick silicone ‘feet’ was added over the tissue, with the intervening space filled with PBS ^20^.

### Quantitative Analysis

Each Swiss-Roll tissue-section was imaged using a mosaic of 8×12 image frames comprising 1200×800 (12-bit) pixels via a cooled digital camera system (Photometrix Quantix) and automated stage (Ludl) connected to a Nikon Fluorescence Microscope (Nikon Eclipse 80i), 20X magnification, all controlled by IPLab software. The light source was an LED based Xylis III (X-Cite). Three sequential color-mosaics were made; 1^st^ using an A488 filter for γH2AX (GH), 2^nd^ a Cy3 filter for anti-EF5 antibody (RS) and 3^rd^ a Cy5 filter for DRAQ5 (DQ)^20^. Subsequently, each image frame had background subtracted via a camera image of PBS, and field-flattening was realized using a custom hemocytometer-based microfluorometer, with appropriate dye standard^20^. All image frames were then converted to 16-bit TIFF files and assembled as a single Mosaic image in MATLAB, for each color (GH, EF and DQ) of each tissue. The mosaics were divided into 4 rows and 4 columns of analysis-squares, each comprising 2×3 image frames (2400^2 pixels) – see **Fig. S1b**.

Each analysis-square triad (e.g. analysis-square in row2, col3 of DQ, GH, and EF stains) was selected based on representative histology, obvious artifacts removed (using ImageJ) and then assessed by a separate MATLAB program (see supplement for details). This program produced a binary tissue/non-tissue mask based on the DQ and GH images and then a 2D array that showed, for each pixel in the mask, its X-Y coordinates, pixel values for DQ, GH and RS, and a normalized value of GH (GH_n_, i.e. 1000xGH/DQ). The factor of 1000 was used to avoid fractional values. The program also calculated a linear best-fit of GH *versus* DQ, and GH_n_ *versus* RS, plus it listed column averages of all parameters, made a histogram of the distributions of each parameter, and made an RGB color image of each analysis square. It should be noted that for any pixel analyzed, one would expect the GH signal to be proportional to the amount of DNA damage present. The proportionality constant is likely to vary with other cell parameters^21^, but represents the best available way to account for the large variation expected in DNA per pixel.

The above-described methods were reproducible enough to allow the use of exposure times for each fluorescence color that remained the same for each tissue of all animals assessed. These times were chosen to encompass the full range of values found, while minimizing saturation of camera pixels. Since the images were 16-bit TIFFs arising from12 bit original images, the range of values for DQ, EF and GH was about 2000 – 65,000, including background.

### Statistical Analysis

The data was further analyzed by calculating the linear regression for each image of each mouse in all dose groups. The slopes and intercepts for all analyzed images (approximately five per mouse, n=3 mice) within each irradiation group were averaged, and statistical significance of these averages between each group was calculated using MATLAB’s two-sample t-test function (ttest2). P-value significance was defined as follows: p > 0.05 – not significant (ns), p < 0.05 – statistically significant (*), p < 0.01 – very statistically significant (**), and p < 0.001 – highly statistically significant (***).

## Results

### EF5 binding in jejunum

The biological half-life of EF5 in mice is about 40 minutes^22^ so most drug had been excreted/metabolized by the time of irradiation (∼2 hours post-injection). Thus, EF5 binding (designated as ‘RS’) represented the average hypoxia level prior to radiation. In contrast, any resulting oxygen-dependence of DNA damage would be reflected in the GH signal, at the time of radiation, and thus in principle, could detect ROD induced decreases in oxygenation during dose delivery. An example of an analysis square for 5 Gy conventional vs zero Gy control is shown in **Fig. 2**. It is visually obvious that the brightest EF signal occurs in the villi tips, with also some fairly bright regions found in the peripheral smooth muscle. In the intermediate crypt region, there was no indication of enhanced EF5 binding. Despite intra-animal variability in overall EF signal, the villi were always found to have the highest levels, muscle intermediate, and crypts lowest values of EF5 binding. Although the subjective interpretation of absolute intensities is difficult in 3-color images, it is quite obvious that the highest GH signal is in regions of low EF and *vice versa* (**Fig. 2, Fig. S2**).Image modification steps for this analysis are detailed in **Fig. S3**.

**Figure 2.**
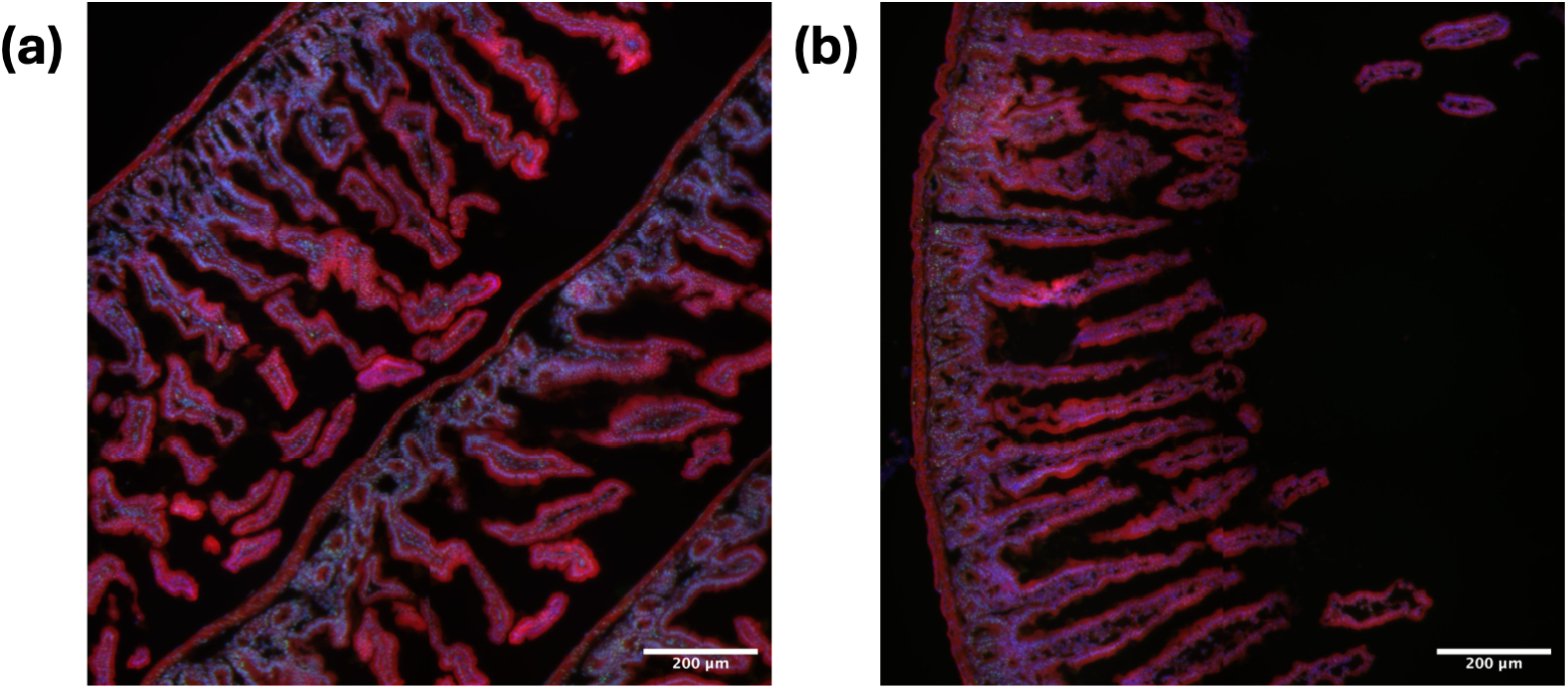
Three-color image (red – EF5, green – GH, blue – DQ) for representative 2×3 field analysis squares for 5 Gy CONV irradiated (a) versus 0 Gy (b) jejunum. One can easily see the substantial decrease in GH (green) for the control tissue (b) and the hypoxic areas (brightest red) of the irradiated tissue (a). Note that these hypoxic areas are dominated by the distal portions of villi and some of the guts peripheral musculature. In order to see the GH signal in the presence of DQ, it has to be enhanced by 3 and EF by 1.25 for both images. This maintains the relationship between irradiated and unirradiated tissue because imaging for all tissues used a constant color-dependent camera exposure for every tissue.

### Hypoxia-dependent DNA damage in jejunum

As detailed in the Supplement, analysis of several (typically 4-6) 2×3 image frames per tissue resulted in about 40-60K tissue-involved pixels each, documenting individual values for DQ, RS, GH and GH_n_. Plots of GH_n_ vs EF for conventional dose-rate showed that damage was inversely dependent on the EF value, suggesting that the degree of hypoxia found was sufficient to allow radioprotection (**Fig. 3**). In animals treated with FLASH dose rates, the slope of this line increased, as predicted for the impact of ROD. This effect was larger at 5 Gy than 2.5 Gy, also expected considering that the amount of ROD is dose dependent. Note that at 5 Gy the entire curve for FLASH fell below that for conventional dose-rate, also supporting an overall decrease in dose-effectiveness for FLASH (FLASH-ntPF mechanism). This was the main effect seen at 2.5 Gy (**Fig. 3**). For unirradiated tissue (**Fig. 2b**) hypoxia only slightly decreased the damage observed at higher levels of hypoxia (see discussion).

**Figure 3.**
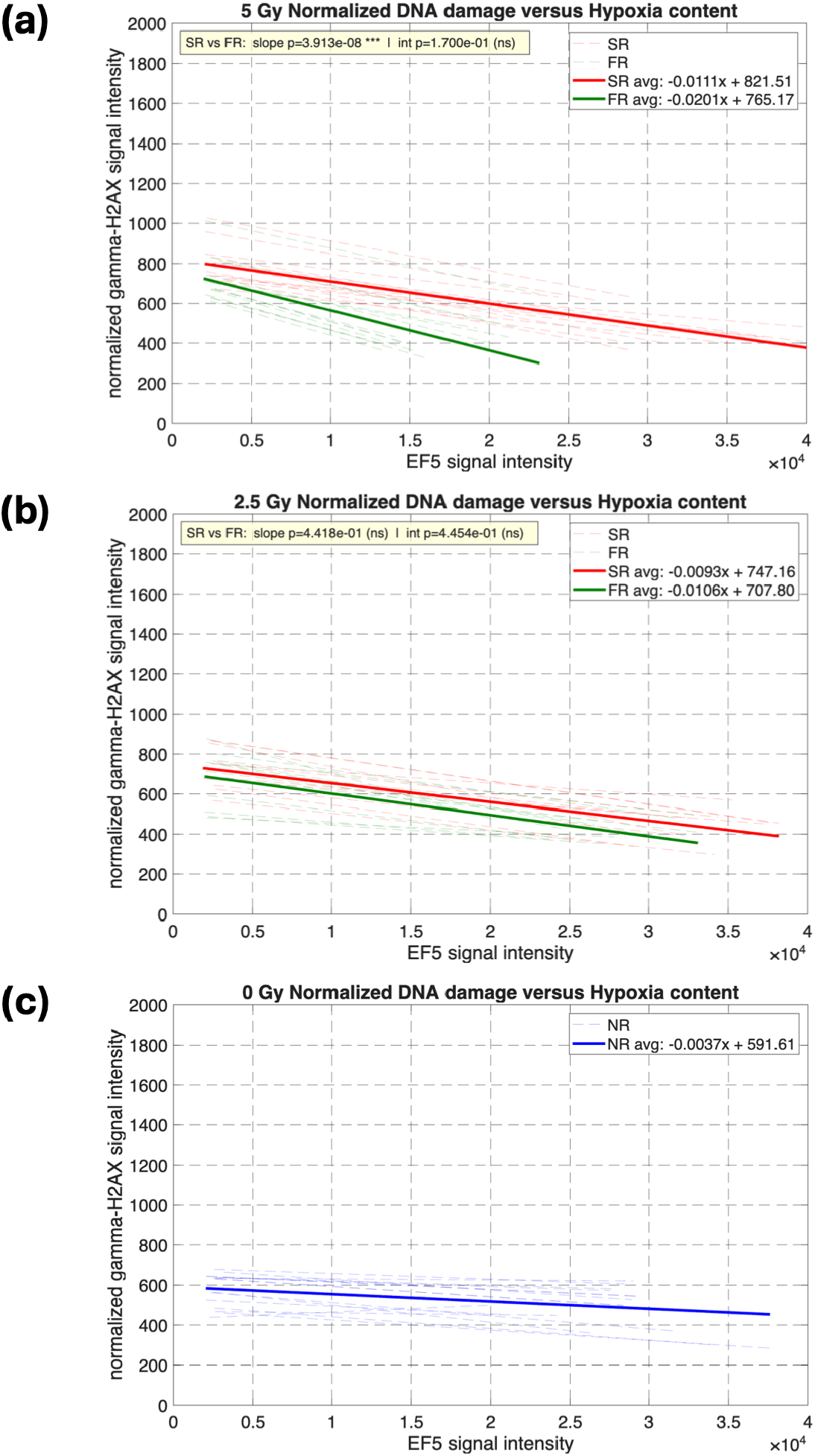
Quantitative evaluation of the relationship between normalized DNA damage (GH_n_) and hypoxia (RS) at 5 Gy (a), 2.5 Gy (b), and 0 Gy (c). Plots (a) and (b) show the conventional dose-rate response in red, and the FLASH dose-rate response in green. At 5 Gy, there is an obvious increase in slope for FLASH as well as an overall decrease in total damage. At 2.5 Gy, the slope change is not significant but there is a trend towards net decrease in damage for FLASH. Controls (c) also show a small negative slope with is under current study.

Turning to the DNA damage signal (GH), we found a strong dependence on DNA content, as suggested above, with lowest slopes for controls and highest for 5 Gy (**Fig. 4**). For the 5 Gy curves there was a clearly enhanced dose-effectiveness observed for conventional vs FLASH dose-rates. Detailed visual inspection determined the pixel-pixel variation in fluorescence was from the totality of anti-γ-H2AX antibodies, not the 3-D variation in number of individual γ-H2AX foci. Thus, as reported previously, we found a sharp plateau, or saturation in this signal, above 5 Gy^20^ using the present methods (see **Fig. S4**). This prevented the investigation of doses above 5 Gy.

**Figure 4.**
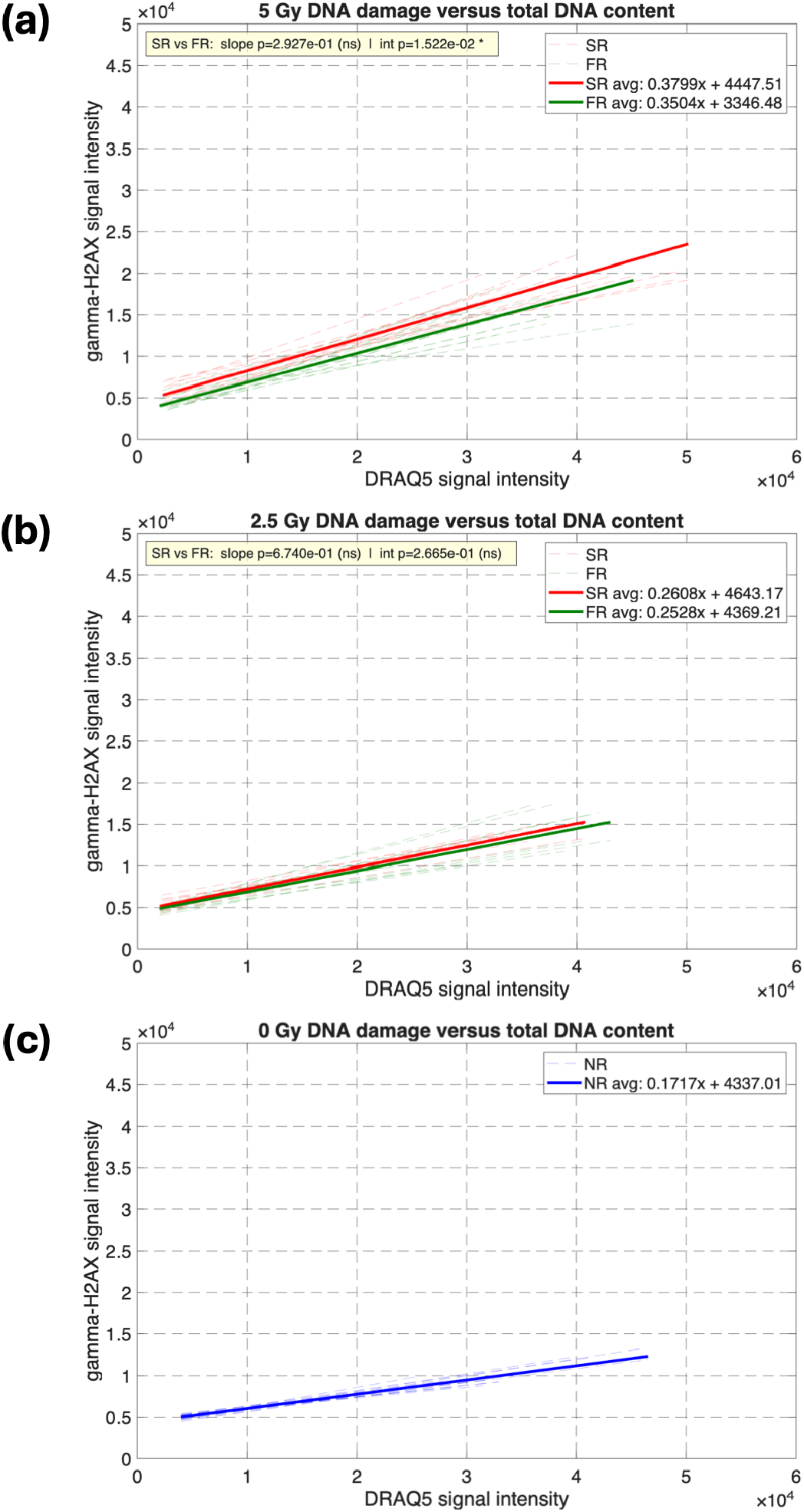
Relationship between GH and DQ as a function of dose at 5 Gy (a) and 2.5 Gy (b). Similar to Figure 2, conventional dose-rate response is shown in red, and the FLASH dose rate response is shown in green. For all tissues, there is an overall increase in DNA damage as total DNA content increases. Controls (c) shows the same relationship (blue). At 5 Gy, although there is not a significant different in slopes, there is an overall decrease in damage in the FLASH irradiated tissues (a).

We used two measures of damage to summarize its overall dose dependence (**Fig. 5**).

**Figure 5.**
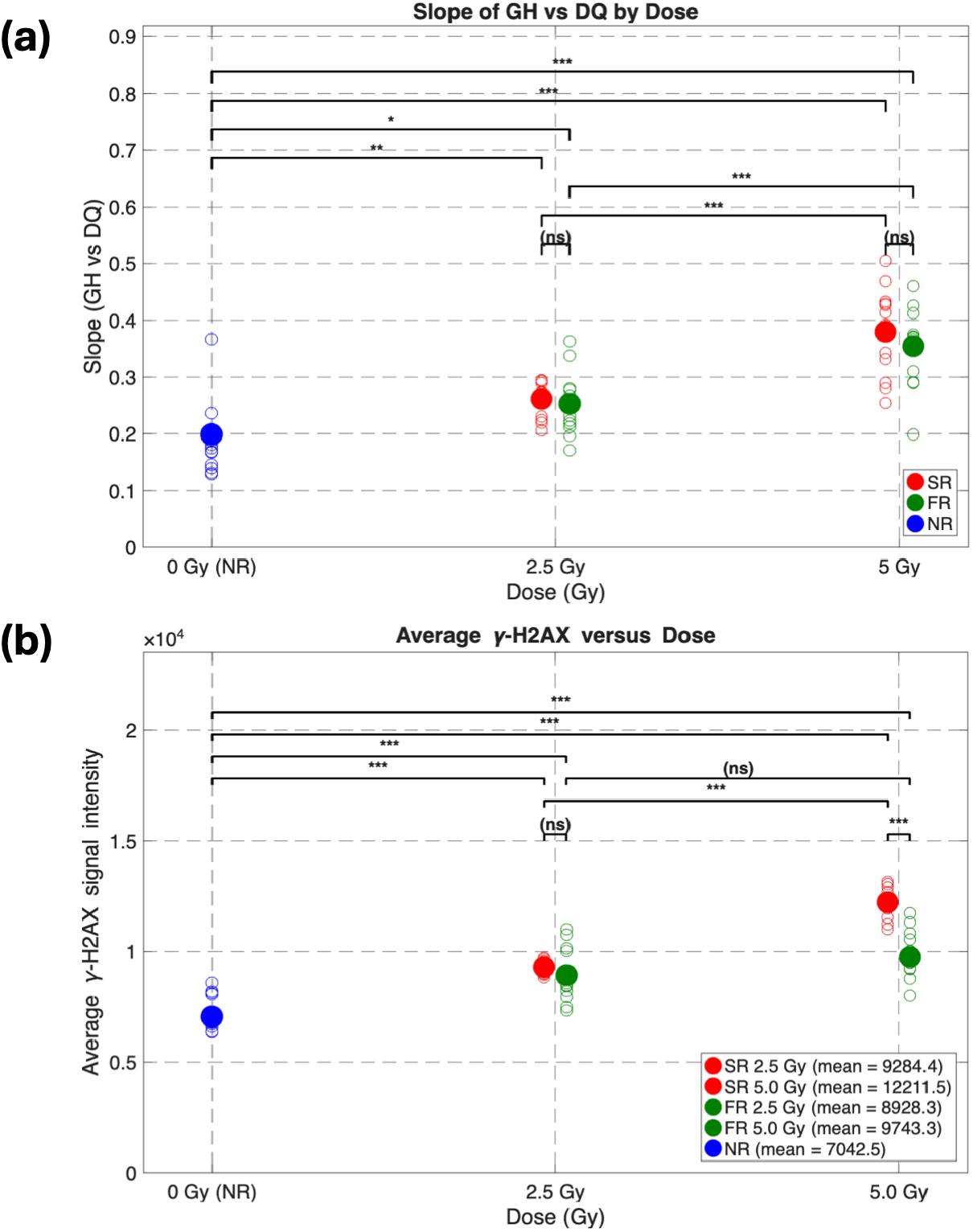
Dose response for the relationship between GH and DQ (a) or average GH (b). Blue represents unirradiated tissues, red represents conventional irradiation, and green represents FLASH irradiation. Levels of significance are shows at top (not significant (ns) – p > 0.05; * – p < 0.05; ** – p < 0.01; *** – p < 0.001). There is a significant increase in slope as dose increases when comparing against the same dose rate (i.e. 2.5 Gy CONV and 5 Gy CONV) (a). As for evaluating average DNA damage versus dose (b), there is a significant difference between FLASH and conventional dose rates at 5 Gy, showing significant sparing, and there is a significant difference in damage when comparing 2.5 Gy and 5 Gy conventional tissues.

The first was to consider the slopes of GH vs DQ (**Fig. 4**) and the second was to simply consider the average GH signal. Both measures showed highly significant changes in GH *versus* dose (**Fig. 5a**) and the second also showed significant decrease in dose-effectiveness by FLASH (compared with conventional) at 5 Gy (**Fig. 5b**).

Since we have shown clearly that the degree of hypoxia in murine jejunum can cause radioprotection, an overall concern might be that the group of FLASH irradiated tumors were by chance more hypoxic (since the degree of hypoxia was a random experimental variable between mice). This was not the case, and in fact, the FLASH irradiated mice were less hypoxic. This was significant for the 2.5 Gy animals, and even more so for the 5 Gy animals (**Fig. 6 – see Fig S5 for examples**). Thus, at constant levels of hypoxia, the decrease in DNA damage allowed by high dose-rate FLASH irradiation would be more substantial than shown in this report.

**Figure 6.**
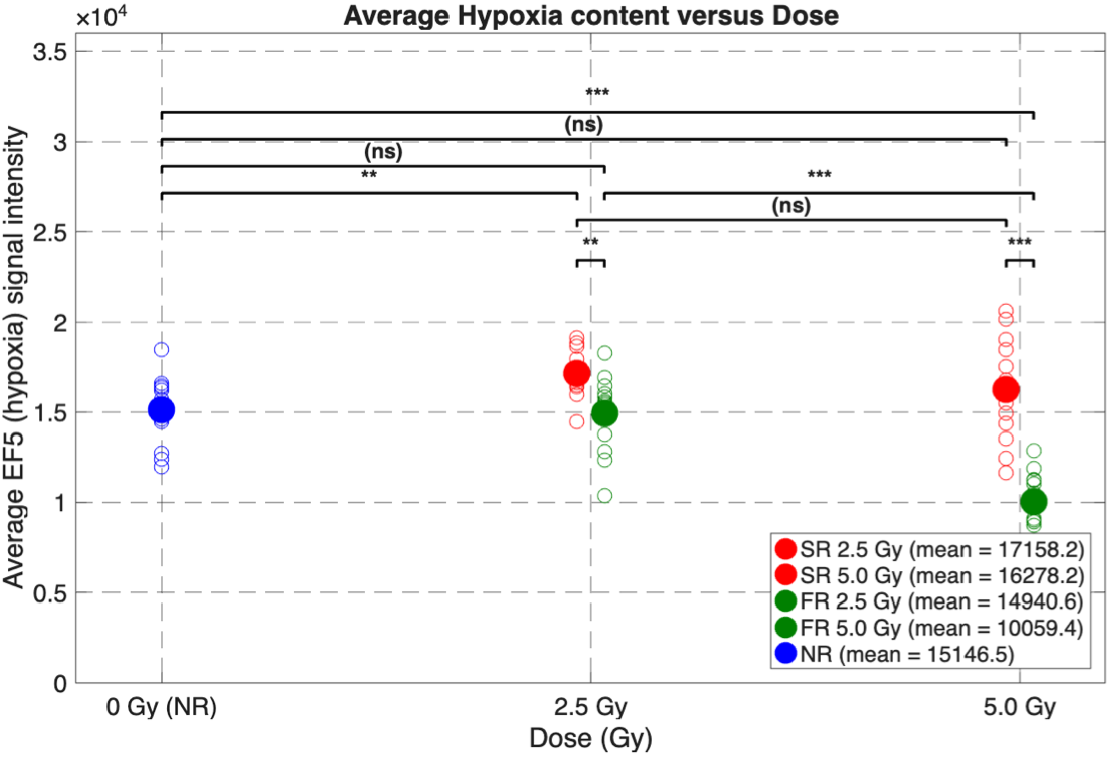
Observed average EF binding level for the 15 mice measured in the imaging experiments, colors as previously described. The FLASH treated groups have decreased EF binding, suggesting higher oxygen levels and greater radiosensitivity, opposite to the measured damage.

## Discussion

The overall goals of this project included documenting, in murine jejunum, the extent and variability of hypoxia, whether observed hypoxia affected the DNA damage observed across the tissue, whether FLASH showed less damage than conventional, and if this damage could have implications for ROD as a significant mechanism underlying FLASH normal tissue sparing. All of these conditions have been observed in this report, and somewhat unexpectedly, we also observed FLASH-ntPF as denoted by an overall decreased dose-effectiveness by FLASH even without considering hypoxia. This effect was first observed in ROD experiments using simple buffers with albumin^10,16^ but was also observed in much more complex solutions^12^. In this latter paper, it was noted that the dose-rate dependence for oxidative chain reactions extended to the high dose-rate realm and for several different substrates, even though first studied at conventional versus extremely low dose rates in purified lipids^23^.

We presented a novel three-component assay that utilizes binding of the 2-nitroimidazole EF5 [2-(2-nitro-1H-imidazol-1-yl)-N-(2,2,3,3,3-pentafluoropropyl)acetamide] ^18^, γ-H2AX binding, and a total DNA assay. EF5 is a metabolic marker for hypoxia, and can measure tissue pO_2_ over a large dynamic range^18^. γ-H2AX is produced at the sites of double strand DNA breaks that arise from radiation. DRAQ5 is a fluorescent dye that binds directly to DS DNA, and was used throughout this study for its high binding affinity^24^. EF5 and γ-H2AX were previously used for the purpose of differentiating hypoxia subtypes in a two-component assay ^20^. The two-component assay was initially developed for whole cells or macroscopic regions of tumor tissue, where the expectation was that very considerable fluctuations in the degree of oxygenation would apply, with accompanying 3-fold differences in radiation dose response. It was not considered sensitive enough to look at relatively small dose changes (e.g. for FLASH perhaps a 10-20% reduction in dose-effectiveness compared with conventional^25^), modest levels of hypoxia and certainly not at the microscopic level considered here (e.g. pixels of size 2.5×2.5 µm^2). The present work has been made possible by multiple refinements to the original methods. These include, in order of importance, a much more stable light source (solid-state LED vs Xenon arc), a 4x higher microscope objective producing 16-fold larger files, and a method to counteract the inherent variability of quantitative DNA staining. These are described thoroughly in the supplement including the resulting dot-blots for paired variables, somewhat akin to those found in flow cytometry – **Fig. S6**.

This assay was developed to address several limitations of the singular measurement of ROD using phosphorescence probes at FLASH and conventional dose rates *in vivo* and *in vitro*^10,16,26^. First, these probes were only able to measure oxygen in the extracellular space *in vivo*, as the probes were unable to pass through the cell membrane. Therefore, any oxygen depletion occurring within the cell at FLASH dose rates could not be measured directly. Further, at conventional dose rates, any oxygen depletion was continuously replenished by the vasculature, therefore one simply can’t measure ROD *in vivo* at conventional dose rates and thus inferences established by simple molecular solutions cannot be compared with *in vivo* data.

There is considerable debate on whether the amount of oxygen depletion observed *in vivo* in the ROD studies was sufficient to be the main cause of sparing at FLASH dose-rates. Pratx and Kapp^27^ found a 30% decrease in radiosensitivity due to ROD at 10 Gy, but only in the range of 3-5 mmHg. This range significantly increased at radiation doses greater than 30 Gy, but single high doses in that range are generally not clinically applicable.

Additionally, for clonogenic sparing in cell survival, they found that at 40 mmHg, there was no difference between FLASH and conventional, but there was a difference at 4 mm Hg, supporting their hypothesis that that cells in normal tissue must be hypoxic on some level in order to receive FLASH sparing^9,27^. More specifically, they suggest the sparing of stem cells in hypoxic niches with oxygen tensions as low as approximately 7 mm Hg are the cause of normal tissue sparing at FLASH dose rates. However, the presence of hypoxic stem cells could not be confirmed in the studies using phosphorescence probes. Of even greater importance is that we have no definitive information on the oxygen dependence of radiation damage for tissues *in vivo*, nor do we know what pO_2_’s exists for these tissues. These limitations strongly motivate the need to directly determine DNA damage and oxygen levels at sub-cellular resolution for the purpose of investigating ROD as a mechanism causing the differential response in FLASH irradiated tissues compared to conventional.

Murine intestine (specifically the jejunum) was used throughout this study for its combination of stem cells and differentiated cells with possibly variable hypoxia content. While it’s been determined that the intestine is more hypoxic than other tissues ^28^, the hypoxia levels within the various sub-structures of the intestine (crypts, stroma, and villi) have not been previously quantified. Using EF5 we were able to determine the relative hypoxia levels for each region of the intestine along with identifying the relationship between DNA damage and hypoxia for all irradiation conditions.

Using pixel-by-pixel analysis in our in-house developed MATLAB program, we were able to calculate the signal intensities throughout each stained image and save them to a dedicated database for that section of tissue. Along with calculating the raw signal intensities, the program calculated the γ-H2AX signal intensity normalized to the total DNA content (DRAQ5 signal intensity) and saved these values (GH_n_) into the spreadsheet. The amount of DNA per pixel varies dramatically throughout a 10 μm-thick section of tissue depending on cell type, stage in the cell cycle, or the fraction of nucleus within a single pixel, etc. Therefore, it was important to normalize GH signal intensity per pixel to DQ signal intensity per pixel.

Using this algorithm, we were able to quantify the difference in DNA damage in FLASH and conventional dose rate irradiated tissues and its relationship to average tissue oxygenation. If the degree of hypoxia was sufficient to partially protect jejunum tissue, and if ROD was sufficient to enhance the radioprotective effect, then we would expect to see a greater impact of pre-treatment hypoxia; thus, the slope of GH_n_ versus hypoxia should become more negative in FLASH irradiated tissues. In contrast, for the second mechanism to potentially have more significance in the FLASH sparing effect (overall decrease in dose-effectiveness across all oxygen levels) then we would expect an overall decrease in DNA damage for FLASH versus conventional irradiation that was not related to tissue pO_2_. We found both of these possibilities to be present in mouse jejunum, but now need to study these parameters for each critical tissue, especially the crypts which contain the stem cells. A previous study showed better stem cell preservation and accelerated crypt recovery after FLASH radiation compared to tissue irradiated at conventional dose rates.^5^ Therefore, a necessary next step would be to quantify DNA damage in the crypts/stem cells to identify whether there is DNA sparing in those populations following FLASH dose rate radiation.

The GH_n_ versus hypoxia plots were used to determine the oxygen dependence of damage (**Fig. 3**). At 5 Gy, we consistently observed an exaggerated oxygen effect (steeper slope) in the FLASH irradiated tissues compared to conventional. However, this steeper slope was not observed in a statistically significant manner for the 2.5 Gy FLASH tissues, however sparing was still observed at all oxygen levels. This suggests a dose dependence to ROD, which has been observed *in vitro* in previous studies^10,26^. However, those studies did not use doses lower than 10 Gy. It is possible that ROD is not significant enough at 2.5 Gy to make a quantitative impact, or that there is a need for an analysis method more specific than pixel-by-pixel analysis to see these differences (i.e. a nuclear analysis method). Additionally, these results were averaged across a very heterogenous tissue – thus further studies are required segmenting the subregions of the intestine to investigate these relationships across different cell types.

The negative slope observed in the GH_n_ versus EF plots for unirradiated controls should be mentioned as well. The initial expectation was that there should be zero slope, however upon qualitative inspection of the IHC images and the quantitative analysis, there is increased GH signal in the area with the lowest EF signal intensities. This area overlaps with the crypts, where highly proliferative stem cells are located. Not considering radiation, or other forms of damage, measurement of γ-H2AX is also cell cycle dependent, with more signal measured in S phase cells^29–31^. Since stem cells are constantly dividing, it is possible that there was more γ-H2AX signal in the crypts area as a result of the cell cycle phase, which was then reflected in the negative slopes shown in **Fig. 3c**. However, further investigation is required to make any conclusive suggestions.

The GH vs DQ plots were used as a measure of overall radiosensitivity, since the DRAQ5 signal intensity is directly correlated to the amount of DNA per pixel in a tissue section. At 2.5 Gy, the FLASH and conventional tissues had similar response (slopes) and baseline (y-intercept) (although the conventional had a slightly increased slope and intercept compared to FLASH, these differences were not statistically significant). At 5 Gy, the FLASH and conventional tissues had similar response to radiation (conventional had a slightly increased slope, but it was not statistically significant). However, the FLASH irradiated tissue started from a lower baseline of damage. Therefore, this indicates that there is FLASH sparing to DNA overall throughout the tissue, but a more specific method of analysis is necessary to investigate these slopes and intercepts in more depth.

It should be noted that the low doses used herein are tolerable by the tissue without whole-body impact, considering typical gut damage assessment^32^. DNA-damage assays in the past have typically suffered from the opposite problem, with very high doses producing lethally irradiated cells. However, it would have been useful to evaluate the effects of substantially higher doses. We observed a pronounced saturation for GH at above 5 Gy, as also reported previously ^20,31^, and believe this is related to saturation of the total available γ-H2AX (see supplement – **Fig. S4**). In our prior report, we suggested that the ability to see ‘all’ of the available γ-H2AX in phosphorylated form was related to the fixation method, but this too requires additional study. To take a more positive view of this limitation, it is unlikely that clinical use of FLASH will employ large single doses, so our present results seem highly relevant to use of FLASH in fractionated doses^6^. It is also possible that other DNA-damage assays pDNA-PK may be useable at higher doses^31^.

## Conclusions

We have identified the hypoxia content in the subregions of murine intestine, which aided in identifying the overall inverse relationship between DNA damage and hypoxia in FLASH and conventional irradiated tissues. We were also able to quantify the difference in DNA damage between FLASH and conventional radiation across a substantial portion of the whole murine intestine using the Swiss roll method. FLASH, compared with conventional radiation appears to provide increased DNA protection in normal tissue, with additional sparing in the low oxygen regions at 5 Gy, the expected result if ROD were rapidly depleting oxygen during FLASH delivery. Separately, FLASH also produced an overall oxygen-independent reduction in DNA damage, consistent with FLASH-ntPF. However, further studies are needed for more thorough investigation of the tissues using nuclear analysis. The study will also benefit from further investigation into the subregions of the intestine separately.

