## supplement for "Decreased Damage for proton FLASH vs Conventional Dose Rates in Mouse Jejunum Shown by Quantitative Assessment of γ-H2AX"

**Unanticipated Issues with overall staining procedure**

**Methods**

In order to compare the DNA damage at FLASH and conventional dose rates and identify if it has a relationship with hypoxia, three assay components are needed. The assay combines EF5 (hypoxia marker), γ-H2AX (identifying double strand DNA breaks), and DRAQ5 (marking total DNA content). DRAQ5 has a high capacity to pass through the cell as well as a high DNA binding efficiency [1], which was verified when a DRAQ5 stained tissue was compared against a Hoechst 33342 stained tissue. In the DRAQ5 stained tissue, there was a more uniform signal intensity along with a brighter signal in the nuclei of the gut's peripheral muscle compared to the Hoechst stained tissue, which was ideal for the purpose of this study. Additionally, its far-red excitation and emission spectrum prevents any overlap between the emission spectrum of anti-EF5 (Cy3) and anti-γ-H2AX (A488) fluorophores [1].

DNA-staining by DRAQ5

We had originally expected that a simple dilution of DRAQ5 from its 5 mM concentrate would allow a consistent DNA stain that could be used as an internal standard for the imaging process (i.e. varying only with the section thickness) – this was essential because the $GH_n$ signal was normalized by dividing GH by the DQ signal. Instead, we found a much greater variation that seemed dependent on the time from dilution from the concentrate. The manufacturer thought that possibly we were seeing microprecipitation, perhaps caused by sub-normal refrigerator temperature levels. This could lead to widely varying times of redissolution. Whether this was the cause, or perhaps some secondary state that involves association between molecules, we found it essential to dilute the stain 24 hours ahead of use, stored out of light in refrigerator, and then small portions brought to RT at least one hour before use, with this time held constant for each slide.

Slide Mounting for Imaging

We had originally planned to mount the tissue using a coverslip and Vectashield Gold – unquestionably, this allows optimal focusing and long-term preservation of the tissue. However, we found that the relationship between the GH and DRAQ5 were modified by the mounting medium and time of contact, possibly caused by its high glycerin content. This caused substantial variability in overall signals for the two DNA stains, so much so that the dose-response data were much less significant. Thus we returned to the original mounting method (using PBS and a (relatively) thick hemocytometer coverslip, as originally described (Koch, 2018).

Imaging

Each imaging session assessed jejunum tissue from one each conventional and FLASH (irradiated) and one unirradiated control animal. The 8x12 image frames were converted from 12 to 16 bits, and stitched together, after background subtraction and field flattening, using MATLAB, to form a MOSAIC, for each tissue and fluorescence color (see Fig S1 showing all 9 mosaics for a single imaging session).

Each of these resulting MOSAICs was divided into 4x4 'analysis squares' (each containing 2x3 image frames – 2400x2400 pixels) – Fig S1b.

The three colors for several of these analysis squares were then assessed, based on representative histology and lack of artifacts (Fig S3, row a). The images from each color were filtered, using a 5x5 averaging filter (to reduce high frequency signal that affects the mask generation) and tissue masks from the DQ and GH images were combined using a threshold assessment (Otsu's method [2]) (Fig S3, row b). Multiplying each tissue by the mask (Fig S3, row c) had very little impact on the observed tissue except for the peripheral musculature, which contained a relatively small nuclear content and hence had sparse DQ or GH signal. Thus, the main impact of the mask operation was to remove non-tissue areas – typically from approximately 5.8x10^6 to ~ 1.2x10^6 pixels. Finally, the image resolution was also reduced by 5 in each dimension to leave an image of 480x480 pixels (Fig S3, row d). This operation left final analysis pixels of size 2.5μmx2.5μm. It did not alter the obvious visual appearance of the tissue in view of the prior filtering operation, since the pixels were still much smaller than cellular nuclei. However, the resolution reduction left a manageable analysis matrix consisting of 40 to 60 thousand non-zero pixels.

Then, values for each color and each assessable pixel, along with the pixel coordinates were saved in an Excel (.xlsx) file. The signal for EF5 was constant for areas considerably larger than each final pixel (2.5x2.5 $\mu m^2$), while it would be expected that the signal from GH would be dependent on the amount of DNA present. Thus, a separate column of data was generated by dividing the GH signal by the DQ signal (GH normalized = $GH_n$). $GH_n$ was scaled by 1000 to allow high integer values, like the other scales, rather than fractions. In order to assess the hypoxia dependence of damage we plotted $GH_n$ *versus* EF, and to assess the overall dose dependence of damage, we plotted GH *versus* DQ. Included in this data matrix were linear regressions of these two plots, column averages for all four parameters, and histograms of each parameter. The program also generated RGB images of the combined parameters.

Saturation of GH at 5 Gy in murine jejunum

Our finding that the anti-γ-H2AX signal saturates at 5 Gy may appear inconsistent with the vast literature on this antigen. We believe that it is related to the superior recovery of this antigen using ethanol vs paraformaldehyde fixation, and was reported previously (Koch 2018), but also recently supported by Kyle and Minchinton[3]. We looked for an increase in detectable signal above 5 Gy in 2 ways: the first was to compare damage observed for 5 Gy vs 10 Gy irradiation using 3-fold higher anti-γ-H2AX antibody. There was essentially no change observed, although non-irradiated tissue showed a somewhat higher background (data not shown). Secondly, we did a full analysis of 5 Gy vs 10 Gy irradiated tissue without changing the staining procedure. It was observed that there was a minor increase for the γ-H2AX signal in the hypoxic portions of the tissue (e.g. villi tips) but no overall change. Analysis showed no change in signal for FLASH vs Conventional, and no oxygen dependence of $GH_n$ vs EF (See Fig S4). Furthermore, the absolute values for the slope of GH vs DQ was the same as observed at 5 Gy – see data in Fig 3 of main paper. This leads to the likelihood that all available H2AX is converted to its phosphorylated form at 5 Gy for this tissue.

### Figures

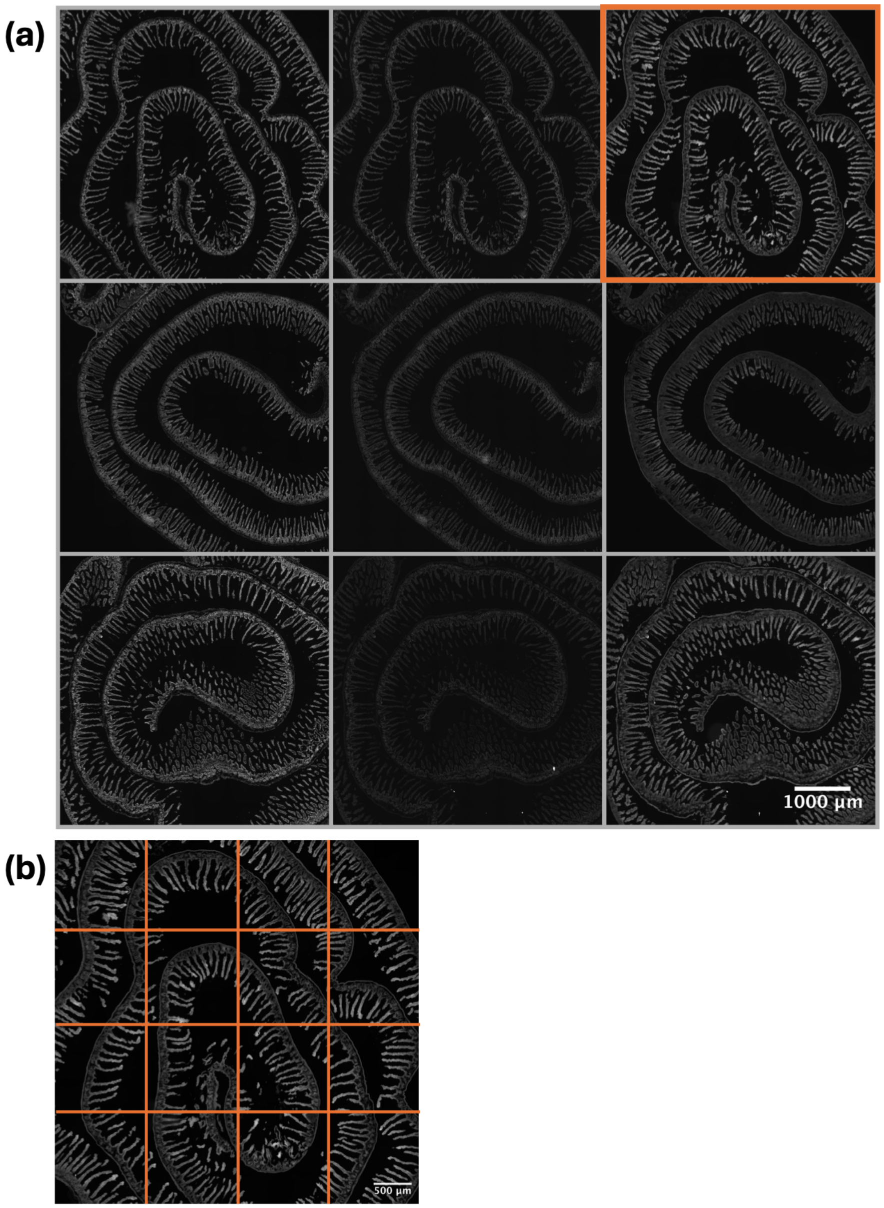


**Fig. S1a. Mosaic of mosaics for one 5Gy conventional (top), 5 Gy FLASH (middle), and control (bottom) tissues.** From left to right of each row, the stains are DRAQ5 (DQ), $\gamma$-H2AX (GH), and EF5 (EF). Note that there is a visual difference between the signal intensities in the GH stain for the conventional, FLASH, and control tissues. Furthermore, there is a difference in the EF signal intensities between the three conditions as well, despite the fact that all images of the same stain employed identical exposures for all tissues studied (GH 1.6", EF 0.75", DQ 0.15" with LED lamp source set at 50% max).

**Fig. S1b. Example of generation, from each mosaic, of 16 square fields, each containing 2x3 of the original imaging fields.** Example shown is from the top right image of (a), 5 Gy conventional, EF.

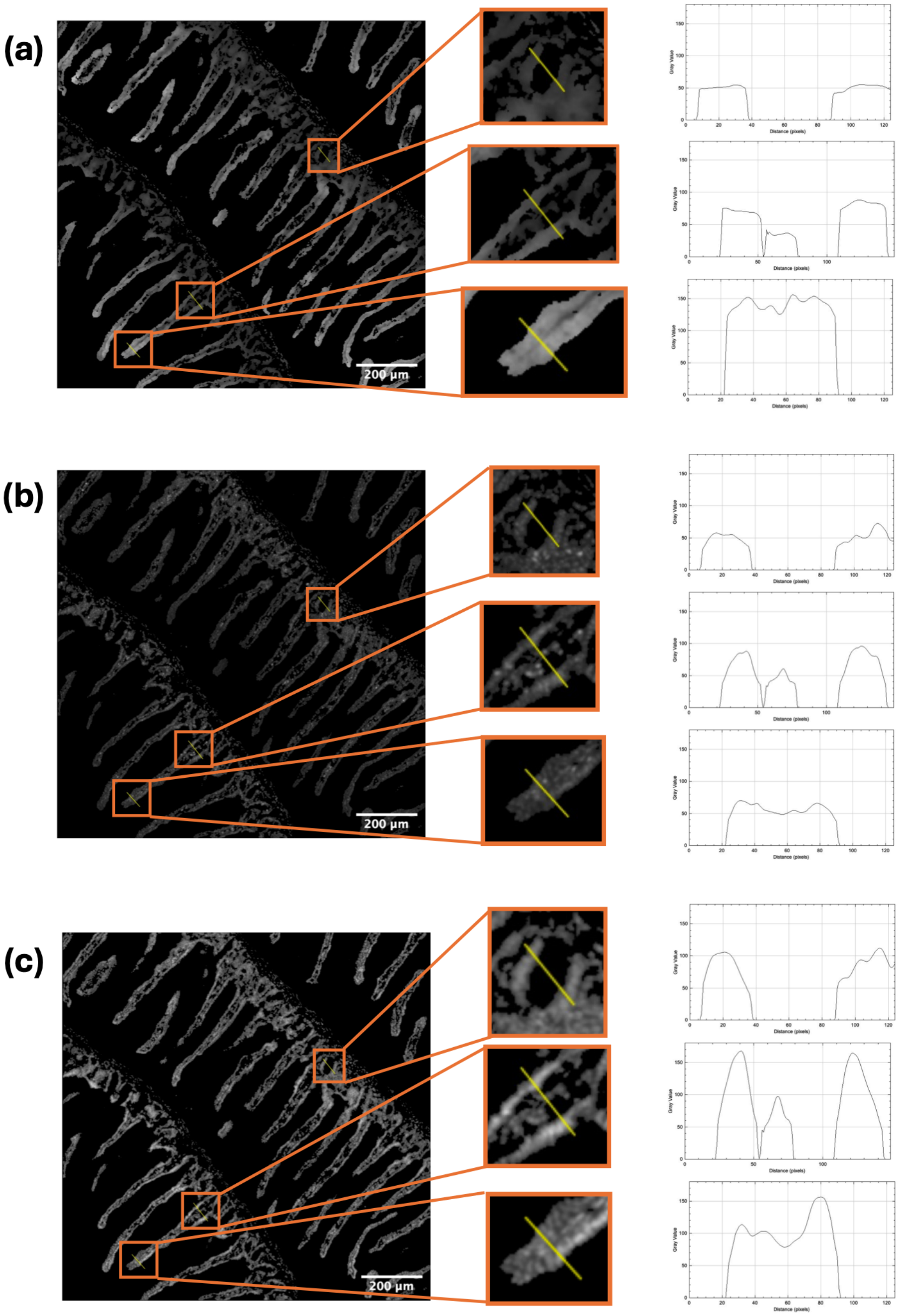


**Fig. S2. Quantification of line segments across villi.** Using one of the 5 Gy tissues (post filtering for analysis), we show that the signal intensities for EF increases from the base of the villi to the tip of the villi (a), and that this is inversed for the GH stain (b) using the Plot Profile feature of ImageJ. Each plot corresponds to the line plot of the zoomed-in image directly to the left of it, and the range of values on the y-axis have been synced for easy comparison.

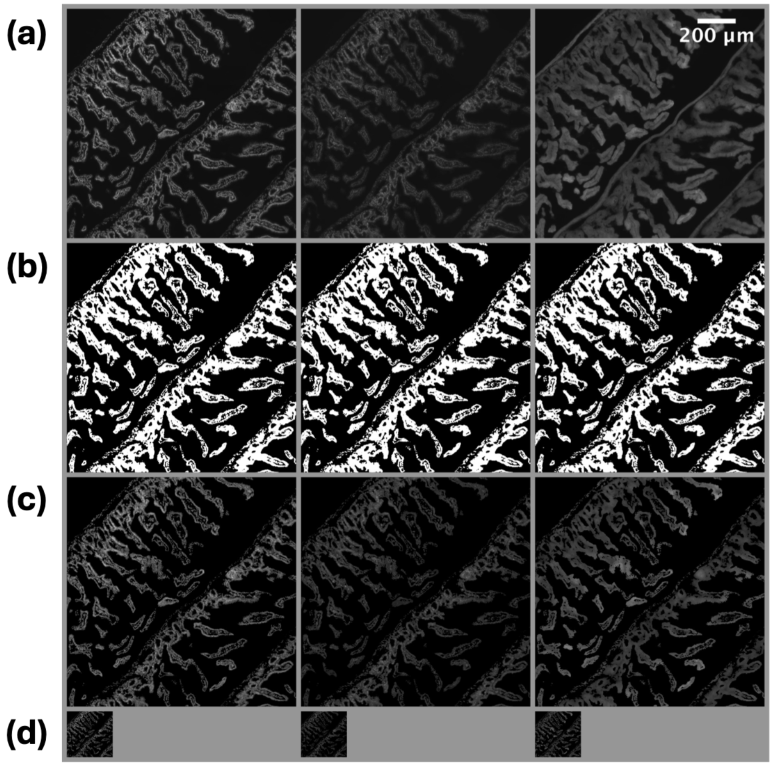


**Fig. S3. Processing tissue data based on the tissue mask from DQ and GH images.** For all rows, the order of stains from left to right is DQ, GH, EF. Row (a) shows the original images before processing. Row (b) shows the binary analysis mask based on the DQ and GH images. Row (c) shows the result of the mask being multiplied to the original images (most noticeable in the peripheral musculature of the EF image). Lastly, row (d) shows the images after increasing the pixel size by 5, thus greatly reducing the total number of pixels for analysis, and hence rows in the analysis matrix (from roughly 10^6 to 40,000).

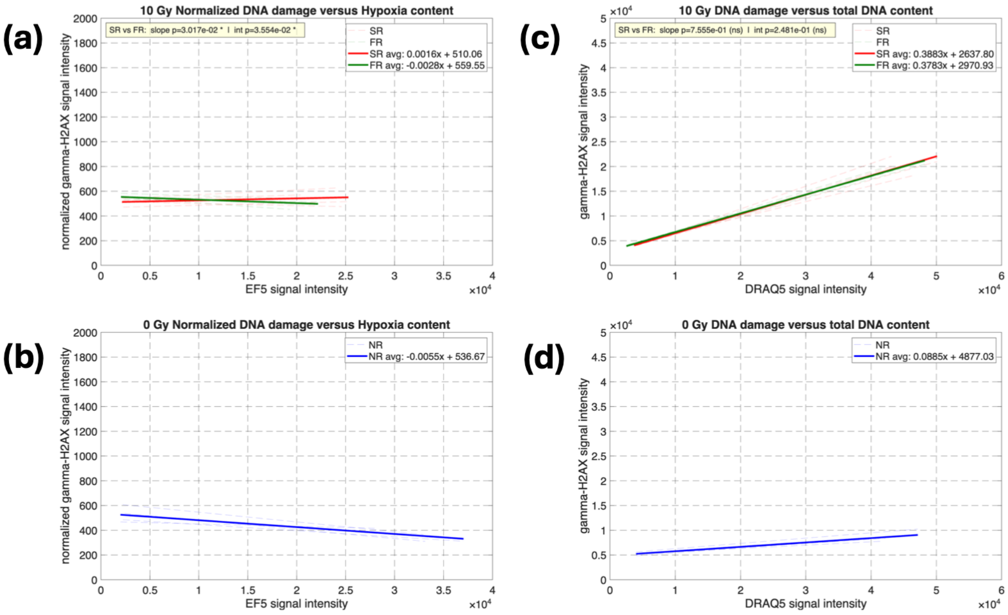


**Fig. S4. Saturation of DNA damage between 5 and 10 Gy.** At 10 Gy, there was a complete loss of the oxygen effect in the irradiated tissue (a). Additionally, the FLASH and conventional linear regressions when plotting GH v DQ completely overlap (c), further demonstrating the saturation of $\gamma$-H2AX at doses between 5 &10 Gy.

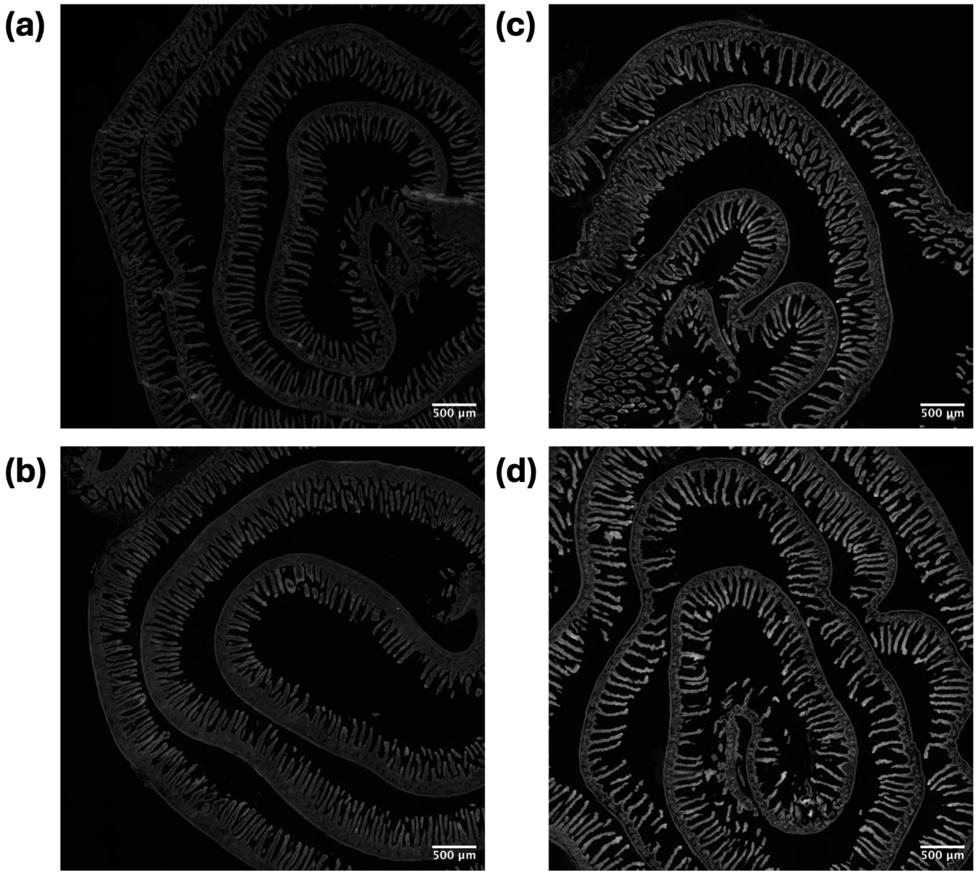


**Fig. S5. Examples of most and least hypoxic tissues of the 5 Gy FLASH (a-b) and conventional (c-d) groups.** As hypoxia increases, the EF5 signal intensity increases proportionally. Therefore, it's clear that the most hypoxic FLASH tissue is less hypoxic than the least hypoxic conventionally irradiated tissue.

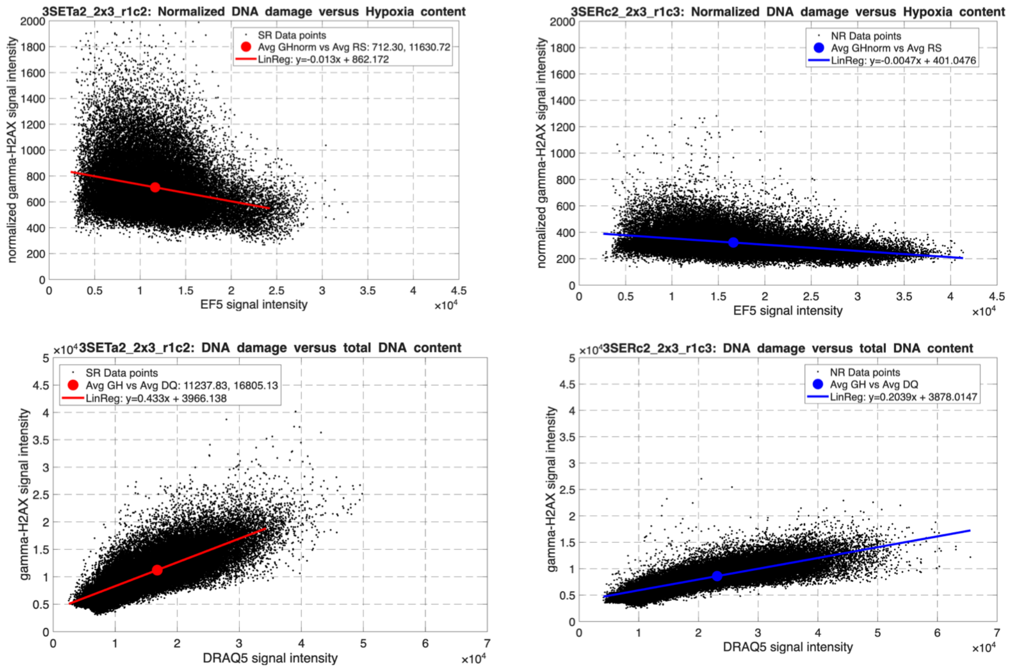


**Fig S6. Dot plot of analysis square for CONV (left column) and Control (right column) tissues ($GH_n$ versus EF – top, GH versus DQ – bottom).** The linear regression for the data is plotted on top of the raw data, and the average of the data is plotted on top of the linear regression. Note that the average of the data consistently goes through the regression line. These plots somewhat resemble 2-parameter flow cytometry images, but the vast number of points somewhat masks the goodness of fit for the linear regressions.
